# *Chlamydia trachomatis* modulates the expression of JAK-STAT signaling components to attenuate the Type II interferon response of epithelial cells

**DOI:** 10.1101/2024.01.09.574898

**Authors:** Francis L. Fontanilla, Rey A. Carabeo, Amanda J. Brinkworth

## Abstract

*Chlamydia trachomatis* has adapted to subvert signaling in epithelial cells to ensure successful intracellular development. Interferon-γ (IFNγ) produced by recruited lymphocytes signals through the JAK/STAT pathway to restrict chlamydial growth in the genital tract. However, during *Chlamydia* infection *in vitro*, addition of IFNγ does not fully induce nuclear localization of its transcription factor STAT1 and target gene, IDO1. We hypothesize that this altered interferon response is a result of *Chlamydia* targeting components of the IFNγ-JAK/STAT pathway. To assess the ability of replicating *Chlamydia* to dampen interferon signaling, HEp2 human epithelial cells were infected with *C. trachomatis* serovar L2 for 24 hours prior to exposure to physiologically relevant levels of IFNγ (500 pg/mL). This novel approach enabled us to observe reduced phospho-activation of both STAT1 and its kinase Janus Kinase 2 (JAK2) in infected cells compared to mock-infected cells. Importantly, basal JAK2 and STAT1 transcript and protein levels were dampened by infection even in the absence of interferon, which could have implications for cytokine signaling beyond IFNγ. Additionally, target genes IRF1, GBP1, APOL3, IDO1, and SOCS1 were not fully induced in response to IFNγ exposure. Infection-dependent decreases in transcript, protein, and phosphoprotein were rescued when *de novo* bacterial protein synthesis was inhibited with chloramphenicol, restoring expression of IFNγ-target genes. Similar *Chlamydia*-dependent dampening of STAT1 and JAK2 transcript levels were observed in infected END1 endocervical cells and in HEp2s infected with *C. trachomatis* serovar D, suggesting a conserved mechanism of dampening the interferon response by reducing the availability of key signaling components.

**Importance:** As an obligate intracellular pathogen that has evolved to infect the genital epithelium, *Chlamydia* has developed strategies to prevent detection and antimicrobial signaling in its host to ensure its survival and spread. A major player in clearing *Chlamydia* infections is the inflammatory cytokine interferon-γ (IFNγ), which is produced by immune cells that are recruited to the site of infection. Reports of IFNγ levels in vaginal and cervical swabs from *Chlamydia*-infected patients range from 1-350 pg/mL, while most *in vitro* studies of the effects of IFNγ on chlamydial growth have used 15-85 fold higher concentrations. By using physiologically-relevant concentrations of IFNγ we were able assess *Chlamydia’s* ability to modulate its signaling. We found that *Chlamydia* decreases the expression of multiple components that are required for inducing gene expression by IFNγ, providing a possible mechanism by which *C. trachomatis* can attenuate the immune response in the female genital tract to cause long-term infections.

## Introduction

Infection of the obligate intracellular pathogen *Chlamydia trachomatis* is the most widespread sexually transmitted disease of bacterial origin in the world (1, 2). Its prevalence is due to its asymptomatic nature, with infected individuals not seeking treatment or unknowingly transmitting the pathogen. Untreated infections can progress to the upper genital tract causing chronic tissue inflammation leading to irreversible sequalae such as pelvic inflammatory disease and tubal factor infertility in women (3, 4, 5). In response to infection, host immune cells infiltrate the infected tissue to clear infection. This cell-mediated immune response is coupled with the production of the proinflammatory cytokine interferon-γ (IFNγ) that induces the expression of the tryptophan-catabolizing enzyme indoleamine-2,3-dioxygenase (IDO1) in epithelial cells (6, 7, 8). With *Chlamydia* being a tryptophan-auxotroph, it is susceptible to depletion by IDO1-mediated catabolism (9). *In vivo* studies showed that IFNy-deficient mice failed to clear or had delayed resolution of urogenital tract infection (10, 11, 12). *In vitro* studies of *Chlamydia*-infected epithelial cells demonstrated growth inhibition and development of enlarged aberrant chlamydial forms when exposed to high concentrations of IFNy throughout infection (13, 14). These studies underlie the effectiveness of IFNγ in resolving chlamydial infections (15). However, genital tract serovars (D and L2) were able to survive and propagate under IFNy treatment during *in vitro* epithelial infection (16). This indicates a potential mechanism that circumvents IFNγ-mediated growth inhibition.

In addition to IDO1, IFNγ is linked to numerous interferon-stimulated genes (ISGs) with diverse mechanisms of inhibiting pathogen growth (17, 18). Their transcriptional response to the cytokine is mediated by the Janus kinase-signal transducer and activator of transcription (JAK-STAT) pathway. Briefly, IFNγ binding to its cognate receptors drives the expression of ISGs via the recruitment and activation of Janus kinases JAK1 and JAK2 in cells. This in turn phosphorylates the transcription factor STAT1 allowing homodimerization and importin-mediated translocation to the nucleus, driving the transcription of genes important for immune functions (19). Because of the strong selective pressure IFNγ imposes on pathogens, the signaling pathway is commonly targeted for attenuation (18). In the case of *Chlamydia* infection of cultured cells, STAT1 translocation to the nucleus, a critical point of regulation, was reported to be reduced via an unknown mechanism (20). In addition to interference by *Chlamydia*, association of STAT1 with transcriptional co-factors can also be targeted for inhibition. The *Toxoplasma gondii* protein TgIST interacts with activated STAT1 dimers in the nucleus preventing recruitment of cofactors necessary for its transcriptional activity following IFNγ stimulation (21). Other anti-IFNγ strategies include proteolytic degradation of IFN receptors and interferon regulatory factors (IRFs) by cysteine protease gingipains from *Porphyromonas gingivalis*, effectively blocking IFN-induced signaling and preventing expression of ISGs (22). Moreover, viral proteins have also been shown to limit the activation of this pathway by STAT degradation and importin sequestration. Viral pathogens inhibit a variety of targets to prevent IFNγ signaling, including the transcription factor STAT1 (18). Vaccina virus produces VH1, which directly dephosphorylates STAT1 to inactivate it, while Japanese Encephalitis Virus, human cytomegalovirus, and Hepatitis B induce phosphatases to retain STAT1 in the cytoplasm (23, 24, 25). Reduced STAT1 nuclear translocation has also been observed due to inhibitory interactions with viral proteins such as VP24 from Ebola virus and Orf6 from SARS-COV-2, which prevent the association of pSTAT1 with karyopherins (26, 27, 28).

Given the conserved targeting of interferon signaling by intracellular pathogens, we hypothesize that *Chlamydia* modulates the JAK-STAT pathway to attenuate the anti-chlamydial effects of IFNγ. Using cervical epithelial HEp2 cells that was validated to respond homogeneously to physiologically relevant levels of IFNγ, we observed reduced expression and activation of both STAT1 and its kinase JAK2 at early stages of infection, with concomitant reduction in expression of ISGs. This attenuation was rescued when de novo chlamydial protein synthesis was inhibited with chloramphenicol (CM), restoring activation and expression of IFNγ-STAT1 target genes. Importantly, these observations were recapitulated in HPV E6/E7-immortalized END1 endocervical cells infected with *C. trachomatis* genital serovar D, suggesting a conserved pathogen mechanism in subverting the host autonomous immune signaling. Taken together, our results point to a pathogen-driven mechanism to dampen the host cell’s interferon response at the transcriptional level. Because the JAK-STAT signaling pathway is shared by other cytokines (e.g., interferon-beta, IL-6, IL-10), *Chlamydia* interference may have a broader impact in the overall epithelial immune response to *Chlamydia* infection.

## Results

### IFNγ-induced activation of STAT1 HEp2 cells is decreased during *Chlamydia* infection

Following the observation that IFNγ-induced activation of STAT1 is attenuated in *Chlamydia*-infected A2EN cells (20), we sought to characterize the basis and significance of this observation in a more physiologically relevant context by lowering the level of exogenously added IFNγ and adjusting the timing of treatment. IFNγ levels found in vaginal secretions and cervical swabs of *Chlamydia*-infected patients range from 1-350 pg/mL (29, 30, 31, 32, 33, 34), which is 15- to 85-fold lower than typical doses used by others for cell culture experiments (16, 20, 35). In addition, because *Chlamydia* is likely not exposed to IFNγ during a primary infection until they have already established their inclusion niche and are actively replicating, we treated cultures at 24 h post-infection (hpi) to better model this situation. Under the modified treatment conditions, we verified that HEp2 cells remained uniformly responsive to the lower dose of IFNγ (Fig. S1C). This homogeneous response enabled observation and quantification of subtle changes in the expression and activation of JAK-STAT signaling factors that occur during infection. We determined the lowest concentration (Fig. S1A) and shortest duration (Fig. S1B) of IFNγ treatment that reliably produced detectable levels of STAT1 activation and ISG production to be 10 U/ml which corresponds to 500 pg/mL. HEp2 cells were treated with 10 U/mL of IFNγ from 15 to 120 minutes to monitor the phosphorylation status of STAT1 (pSTAT1), and the induction of expression of IDO1 (Fig. S1B). After 2 hours of treatment, there was an apparent activation of STAT1, but no concomitant expression of IDO1 (Fig. S1B). We determined a 6-h IFNγ treatment to be sufficient in activating STAT1 and inducing IDO1 expression at both the mRNA and protein levels (Fig. S1C-E).

To test if actively replicating *Chlamydia* can attenuate the IFNγ response, we infected HEp2 cells with *C. trachomatis* serovar L2 for 24 hours and exposed infected cells to 10 U/mL of IFNγ for another 6 hours. Several indicators of STAT1 activation were monitored. This included quantification of STAT1 nuclear translocation by immunofluorescence confocal microscopy, STAT1 phosphorylation by Western blot, and IDO1 transcript levels by RT-qPCR and protein levels by Western blot. Similar to previous findings in A2EN cells, we observed that there is a significant decrease in pSTAT1 fluorescence intensity in the nucleus of infected cells following IFNγ exposure compared to uninfected cells (Fig. 1A & S1C). This was in agreement with apparent decreased levels of pSTAT1 (Fig. 1C). Importantly, expression of the STAT1-regulated IDO1 gene was also decreased (Fig. 1C & 1D), indicating that the magnitude of reduction of STAT1 activation induced by *Chlamydia* was sufficient to affect expression of a potent anti-chlamydial ISG.

**Figure 1.**
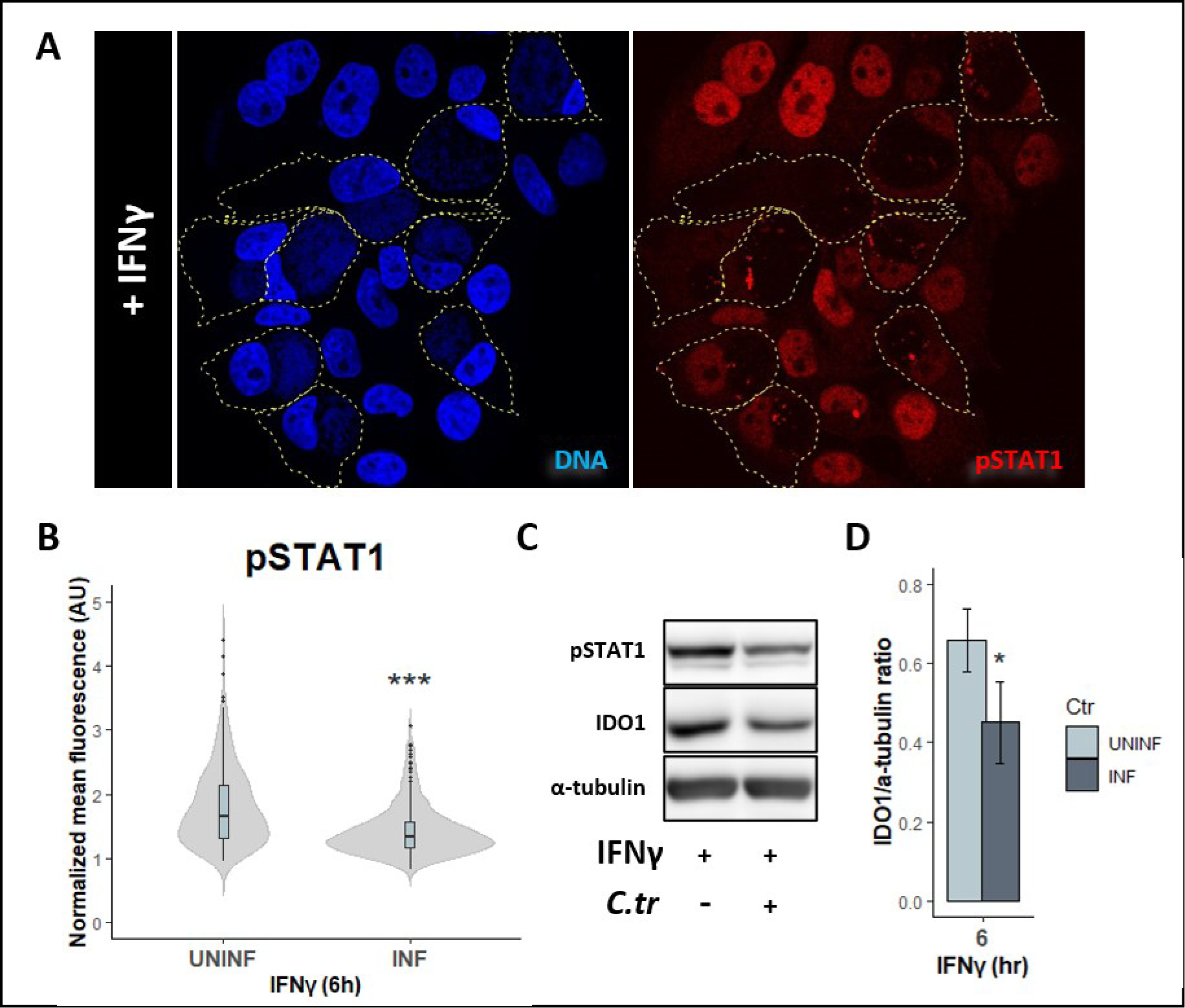
IFNγ-induced activation of STAT1 is attenuated in *C. trachomatis* L2-infected Hep2 cells. **(A)** *C. trachomatis*-infected cells (MOI=2, yellow outline) and uninfected HEp2 cells exposed to IFNγ (10 U/mL) starting at 24 hpi for 6 hours were fixed and stained for phosphorylated STAT1 (pSTAT1) in red and for DNA with DAPI in blue. **(B)** Nuclear pSTAT1 fluorescence intensities were quantified in both uninfected and infected cells. Measurements were normalized to respective no IFNγ control to account for background pSTAT1 staining and plotted as mean fluorescence. Violin plot represents 6 independent experiments with total of 260 nuclei measured per group. Statistical significance was determined by Wilcoxon Rank-sum. **(C)** pSTAT1 and IDO1 protein levels were assessed from total protein lysates collected from *C. trachomatis*-infected (MOI=4) and uninfected cells exposed to IFNγ (10 U/mL) at 24 hpi for 6 hours. **(D)** Densitometric quantification of IDO1 plotted as ratios to the loading control α-tubulin. Bar graph represents 4 independent experiments, and statistical significance was determined using Welch’s t-test. *p < 0.05, ***p < 0.001.

### JAK-STAT signaling pathway components are reduced during infection

The decrease in IFNγ-induced activation of STAT1 could be due to lower expression levels of individual pathway components (e.g., receptors, kinases, STAT1, karyopherins) or to reduced kinase activity. We quantified by RT-qPCR the transcript levels of JAK-STAT pathway genes in IFNγ-exposed mock- and *Chlamydia*-infected cells and observed in the latter reduced transcript levels of IFNGR1, JAK1, JAK2, KPNA1, KPNB1, and STAT1, while IFNGR2 was slightly increased (Fig. 2A). Since JAK and STAT proteins undergo post-translational phosphorylation in response to cytokines, we tested the activation state of JAK2 and STAT1 by Western blotting. We observed lower levels of pJAK2 and pSTAT1 in infected cells compared to uninfected cells following IFNγ treatment (Fig. 2B), confirming that the reduction in transcription produced detectable decreases in phospho-activated JAK2 and STAT1 (Fig. 2C & 2D). Results indicate JAK2 and STAT1 transcription being targeted by the pathogen. Because we were not able to detect JAK1 protein under any conditions by immunoblotting (data not shown), we were unable to confirm if the decrease in transcript levels resulted in decreased protein levels.

**Figure 2.**
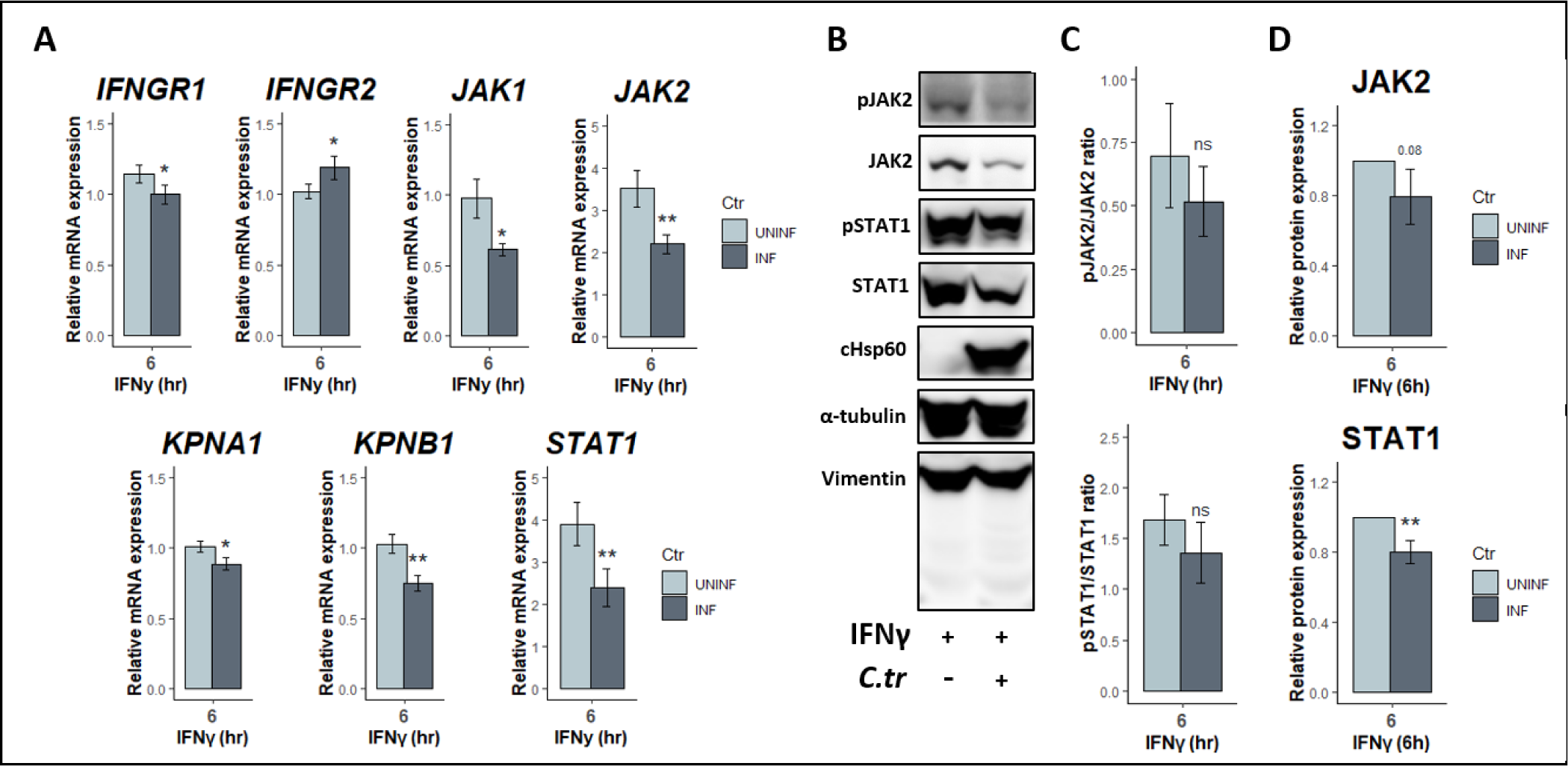
Total JAK2 and STAT1 are transcriptionally down-regulated in infected cells. **(A)** RT-qPCR analysis of JAK-STAT pathway components that are required for IFNγ signaling. ΔΔCTs were calculated by comparing to the mock-infected, untreated sampel and normalized to the housekeeping gene HPRT. All data are representative of 4 independent experiments, and statistical significance was determined using Welch’s t-test. **(B)** Total protein lysates collected from *C. trachomatis*-infected (MOI=4) and uninfected cells exposed to IFNγ (10 U/mL) at 24 hpi for 6 hours were probed for total JAK2 and STAT1 as well as their phosphorylated species. α-tubulin serves as loading control and vimentin was probed to show that there is no infection-induced general protein cleavage during sample collection. **(C)** Densitometric quantification of pJAK2 and pSTAT1 expressed as ratio to their respective total proteins, and **(D)** relative protein expression levels of total JAK2 and STAT1 normalized to the loading control α-tubulin**p < 0.01, ns = not significant.

Our observations show that *Chlamydia* might be targeting the expression levels of these proteins to limit the response of the host following cytokine stimulation. For this mechanism to be effective, we reasoned that *Chlamydia* needs to modulate these pathway components early in infection prior to cytokine stimulation, i.e. preconditioning of host cells by the pathogen. To address this, we monitored the basal expression levels of JAK2 and STAT1 across multiple timepoints in infected, but unstimulated cells. We observed an infection-dependent decrease in STAT1 transcript as early as 8 to 16 hpi in infected cells, and a decrease in JAK2 between 16 and 24 hpi (Fig. 3A & 3B). Reduction in transcripts corresponded with decreased levels of total JAK2 and STAT1 proteins (Fig. 3C). Taken together, our data suggests that *Chlamydia* modulates the levels of JAK-STAT pathway components at the transcriptional level leading to reduced activation by IFNγ (Fig. 2). Since the effects of IFNy on chlamydial development have been shown to be cell line-dependent and serovar-specific (16, 35), we found a similar transcriptional downregulation of these components with *C. trachomatis* serovar D infection of HEp2 cells (Fig. S2) and when infecting the human endocervical cell line END1 with serovar L2 (Fig. S3).

**Figure 3.**
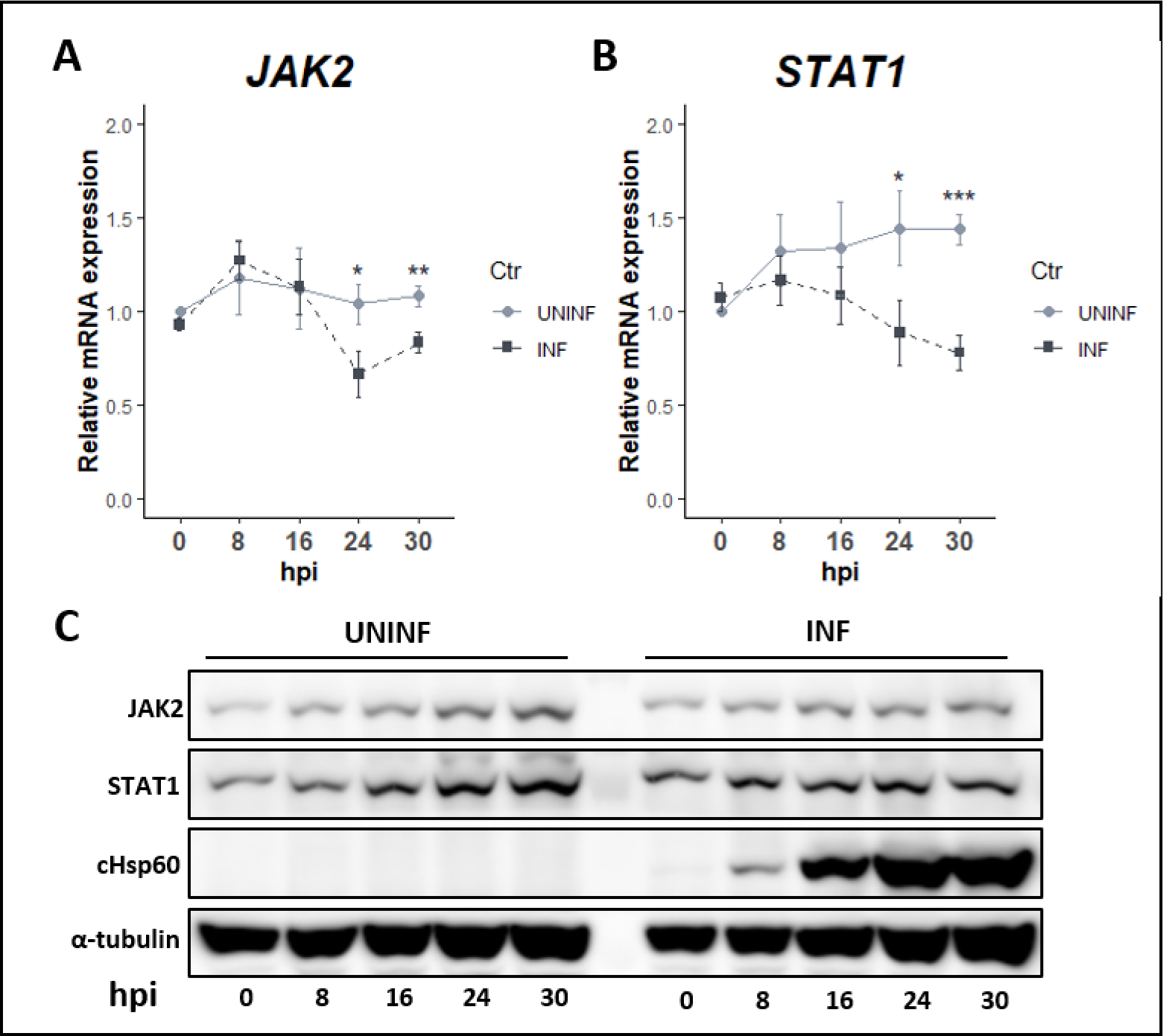
Infected cells have lower basal expression levels of JAK2 and STAT1. **(A-B)** Total RNA was collected from *C. trachomatis*-infected (MOI=4) and uninfected cells at indicated timepoints post-infection, and transcript levels of **(A)** JAK2, and **(B)** STAT1 were quantified by RT-qPCR, normalizing to the housekeeping gene HPRT. **(C)** Parallel protein levels of total JAK2 and STAT1 were also assessed by immunoblotting with cHsp60 as marker for infected cells, and α-tubulin as loading control. All data are representative of 3 independent experiments, and statistical significance was determined using Welch’s t-test. **p < 0.01, *p < 0.05.

### Attenuation in JAK-STAT signaling components is pathogen-driven

During active infection, *Chlamydia* uses effector proteins to hijack and manipulate host cellular pathways to establish an intracellular niche and allow its successful development (36, 37, 38, 39, 40). Our results showing lower JAK2 and STAT1 expression levels during infection prompted us to determine if this required chlamydial protein synthesis. Infected HEp2 cells were treated with the bacterial protein synthesis inhibitor chloramphenicol (CM). This antibiotic binds specifically to 23S rRNA of the prokaryotic 50S ribosomal subunit thereby inhibiting the growth of the polypeptide chain (41, 42). Chloramphenicol (50 μg/mL) was added to infected cells at 8 hpi, since we observed that there is already a decrease in JAK-STAT components at this timepoint (Fig. 3A-B). Treatment with CM was maintained for the remainder of the experiment to 30 hpi (Fig. 4A). Inhibition of chlamydial protein synthesis is reflected by the lower chlamydial cHsp60 protein band intensity and reduced size of chlamydial inclusions in cultures (Fig. 4A, S4). Following CM treatment, we observed that transcript and protein expression of both JAK2 and STAT1 were rescued in infected cells (Fig. 4B-C). Moreover, this also restored the IFNγ-induced phosphorylation of STAT1 but not JAK2 (Fig. 4A). These results suggest that chlamydial *de novo* protein synthesis is necessary for regulating the expression levels of JAK-STAT components, and, hence, the attenuation in IFNγ-induced STAT1 activation.

**Figure 4.**
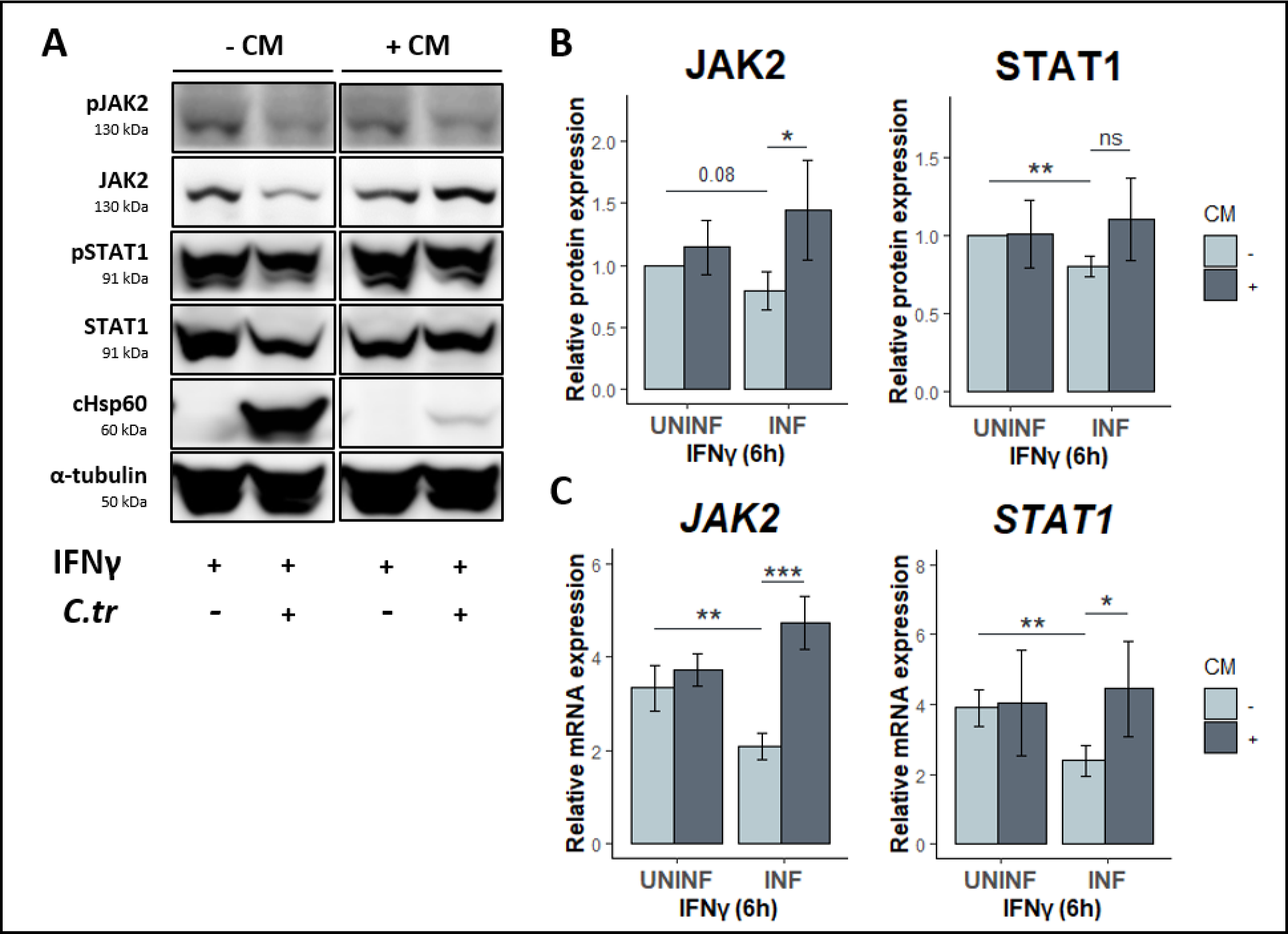
*Chlamydia* protein synthesis is required to down-modulate JAK2 and STAT1. **(A)** To determine if the attenuated phenotype Is pathogen-driven, we used the bacterial protein synthesis inhibitor chloramphenicol (CM) early in the infection at 8 hpi to inhibit the *de novo* production of chlamydial proteins. Total protein lysates collected from infected (MOI=4) and uninfected cells exposed to IFNγ (10 U/mL) at 24 hpi for 6 hours were probed for total JAK2 and STAT1, as well as their phosphorylated species. **(B)** Corresponding densitometric quantification of JAK2 and STAT1 protein levels normalized to the loading control α-tubulin. **(C)** Parallel JAK2 and STAT1 transcripts normalized to HPRT1 were quantified using qPCR. All data are representative of 4 independent experiments, and statistical significance was determined using Welch’s t-test. ***p < 0.001, **p < 0.01, *p < 0.05, ns = not significant.

### *Chlamydia* suppresses expression of IFNγ-target genes

IFNγ activates a plethora of ISGs that are vital in the host immune response against intracellular pathogens (19). The enzyme IDO1 is one of these ISGs and is widely considered an anti-chlamydial host effector through its ability to degrade intracellular tryptophan, limiting the access of this essential nutrient to the Trp auxotrophic *Chlamydia*. Given that *Chlamydia* can target IFNγ signaling for attenuation, we hypothesized that other ISGs are down-modulated as well, and inhibiting bacterial protein synthesis would rescue this down-modulation. We monitored the expression levels of IFNγ targets GBP1, APOL3, IRF1, and IDO1 using RT-qPCR. We quantified a general attenuation in the transcript levels of these genes following IFNγ treatment, and a corresponding increase in transcripts when signaling was restored in CM-treated infected cells (Fig. 5A-D). Overall, our findings suggest that *Chlamydia* is actively down-regulating expression levels of IFNγ signaling pathway components to dampen the host cell response, in concert with its other immune evasion strategies.

**Figure 5.**
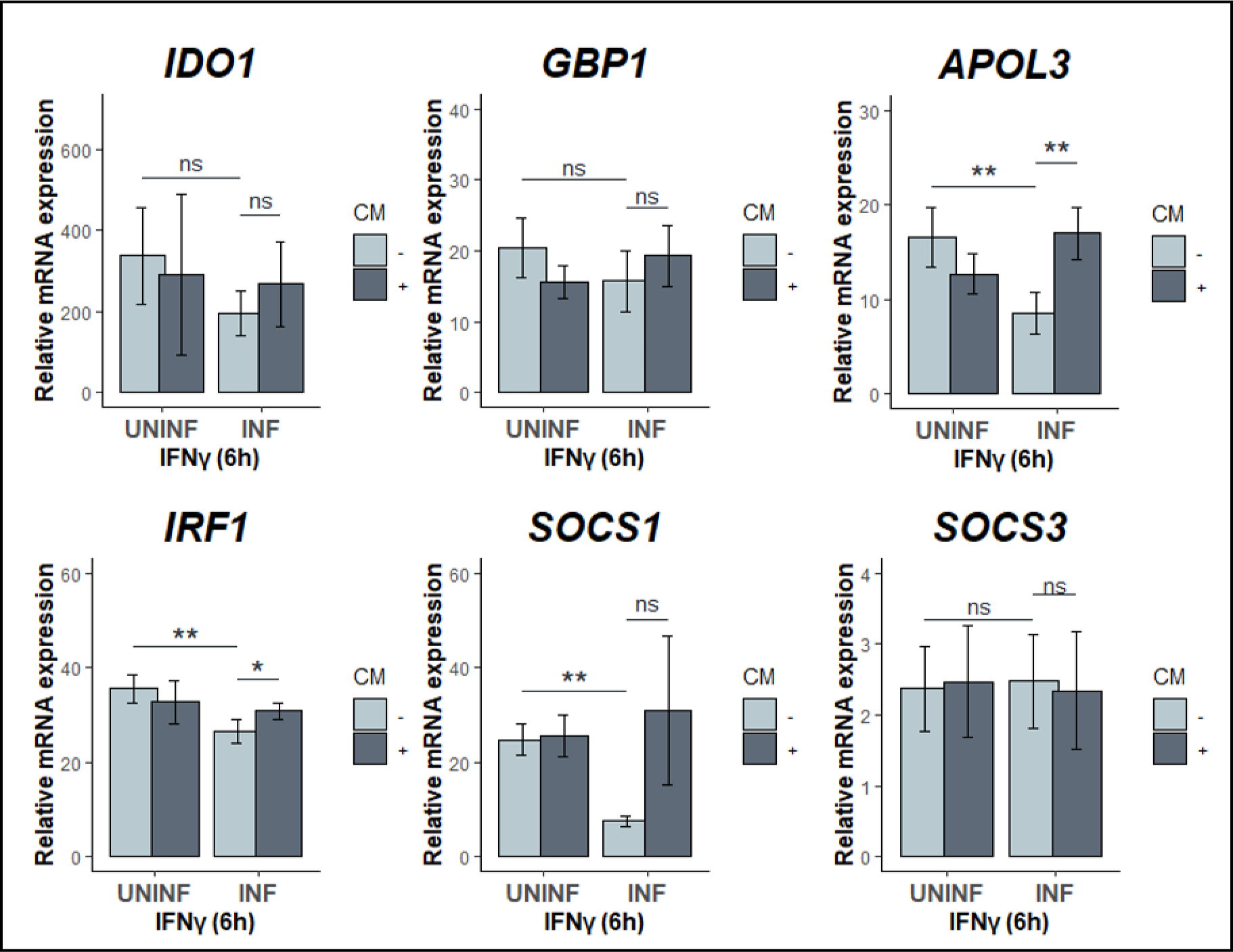
*C. trachomatis* infection drives dampening of ISG expression during IFNγ-exposure. Total RNA was collected from infected (MOI=4) and uninfected cells exposed to IFNγ (10 U/mL) at 24 hpi for 6 hours, with and without chloramphenicol (CM) added at 8 hpi. Transcript levels of IFNγ-STAT1-target genes IDO1, GBP1, APOL3, IRF1, SOCS1, and SOCS3 were quantified by RT-qPCR. All data are representative of at least 3 independent experiments, and statistical significance was determined using Welch’s t-test. **p < 0.01, *p < 0.05, ns = not significant.

## Discussion

Successful pathogens have evolved mechanisms to mitigate host immune responses to aid in their survival and spread. Circumventing the growth inhibitory effect of IFNγ is essential to chlamydial survival in the genital tract. Here, we show that *Chlamydia* targets the expression of key JAK-STAT pathway components at the transcriptional level to dampen the IFNy response of infected epithelial cells. This downregulation happens early in infection, priming cells to respond sub-optimally to IFNy stimulation. While we confirmed that downregulation of JAK2 and STAT1 was consistent at the protein level and activation state, downregulation of other factors such as karyopherins (KPNA/B) and interferon gamma receptor 1 (IFNGR1) may also contribute to dampened IFNγ signaling as shown by the downregulated transcript expression.

Our data indicate that the transcriptional downmodulation of JAK-STAT signaling components requires chlamydial protein synthesis, which suggests that *Chlamydia* produces effector proteins to influence host transcription in a direct or indirect manner. This observation is notable because it has previously been shown that subversion of cellular processes in infected cells requires direct protein-protein interaction with chlamydial type III secreted effectors or inclusion (Inc) membrane proteins (43, 44). While chlamydial effectors have been implicated in relieving the effects of some ISGs and NFκB activity modulation, their roles in regulating the JAK-STAT signaling pathway in epithelial cells remains unknown (45, 46). Although there is a significant increase in transcript levels of the known JAK-STAT negative regulator SOCS3 during infection, which can lead to JAK2 protein degradation, this does not explain the decrease in levels of both JAK2 and STAT1 at the transcript level. Moreover, the IFNγ target SOCS1 follows the attenuated expression pattern similar to other downstream ISGs, suggesting that the dampened IFNγ response affects a wide range of IFNγ-inducible ISGs. We also found that this down-modulation of JAK2 and STAT1 expression during IFNγ exposure is also consistent in genital serovar D-infected HEp2 cells and in serovar L2-infected endocervical END1 cells, indicating that this regulation is serovar- and host cell-independent. Previous studies have demonstrated that expression of STAT1 is induced following *Chlamydia* infection (47, 48). Without use of a protein synthesis inhibitor, such as chloramphenicol, such studies may have missed any limiting effects of chlamydial proteins on epithelial responses to infection. Our chloramphenicol studies suggest that IFNγ signaling is less than optimal following cytokine stimulation due to a *Chlamydia*-driven suppression of JAK-STAT pathway components resulting in suppressed ISG transcription. In addition, it has also been shown that bystander uninfected cells and infected cells respond differently when exposed to IFNγ (20), further suggesting that analyzing JAK-STAT signaling components at a population level requires a monolayer of fully infected cells which might not be recapitulated when using lower MOIs.

Our study detected a marginal, but consistent, decrease in downstream ISGs in IFNγ-exposed infected cells, suggesting that *Chlamydia* not only limits the expression of IDO1, but other relevant ISGs with confirmed or suspected anti-chlamydial effects. The IFNγ target GBP1 has been shown to impact C*hlamydia* development. GBPs are GTPases that bind to LPS of a variety of Gram-negative pathogens, resulting in the disruption of bacterial cell envelope (49, 50, 51). In the context of *Chlamydia* infection, chlamydial inclusion size was observed to be smaller in cells when GBPs are exogenously expressed (52). The mechanistic basis for the anti-chlamydial function of GBPs in *Chlamydia* has been reported to require an intact GTPase activity (52). Moreover, GBP1 has been implicated in the activation of inflammasome and cell death in *Chlamydia*-infected cells, further suggesting a potential anti-chlamydial role (53). The ISG APOL3 has been shown to work in tandem with GBP1 in targeting and restricting the growth of several cytosol-invasive bacteria (54). The exact function of APOL3 remains unknown, and it has not yet been investigated in *Chlamydia*-infected cells. In short, expression of several ISGs are induced by IFNγ, and whether they are all relevant to *Chlamydia* infection needs to be addressed, with multiple ISGs acting in combination to control chlamydial growth. Their potential synergism with other ISGs further underscore the need for *Chlamydia* to regulate the IFNγ response in infected cells. Whether the magnitude of attenuation of IFNγ by *Chlamydia* is sufficient is not known.

It is also possible that attenuation of the JAK-STAT signaling pathway may be more relevant in the context of other cytokines. Numerous studies have demonstrated that it takes at least 24 hours of IFNγ pre-treatment to affect chlamydial growth (16), so it is still unclear how down-modulation of these signaling components may help *Chlamydia* during primary infection. However, this regulation of JAK-STAT signaling components in infected cells may have broader implications on how infected epithelial cells respond to cytokines. The JAK-STAT pathway serves as a platform for other relevant interferons and cytokines for signal transduction. Previous studies have shown that infected epithelial cells produce a plethora of cytokines that may act both in an autocrine and paracrine manner (48, 55, 56). One of these cytokines is the closely related interferon-β (IFNβ) which is important in the general cell-autonomous immune response of epithelial cells against intracellular pathogens, particularly during initial stages of infection (57). We surmise that dysregulation of the JAK-STAT signaling components early in infection may have negative consequences on the IFNβ autocrine response of infected cells preventing the induction of an intracellular antimicrobial state. In addition, both signaling and production of interleukin-6 (IL-6) are dependent on the JAK-STAT pathway (58). This cytokine is important in immune regulation as it promotes recruitment and activation of immune cells following pathogen infection (59). Thus, perturbation in the IL-6-JAK-STAT axis may lead to altered IL-6 signaling outcomes such as inadequate recruitment of effector immune cells to facilitate pathogen restriction. Indeed, intact IL-6 signaling is required in clearing *C. muridarum* infection in mouse genital tract (60). Further study is required to determine the overall consequence of modulation of JAK-STAT signaling in *Chlamydia*-infected epithelial cells on immune cell recruitment and activation, as well as, on potential effects on neighboring epithelial cells to support secondary chlamydial infection. An active epithelial cell-based innate immune response could prompt acquisition by *Chlamydia* of a JAK-STAT attenuating mechanism through evolution. Epithelial cells having an active role in limiting infection could be more relevant to secondary infection dynamics, where timing of infection and exposure to cytokines become significant factors.

## Materials and Methods

### Cell line culture conditions

Human cervical carcinoma cell line HEp2 cells (ATCC® CCL-23) were cultured at 37°C with 5% atmospheric CO_2_ in Dulbecco’s Modified Eagle’s Medium (DMEM; Gibco, Thermo Fisher Scientific, Waltham, MA, USA) supplemented with 10% (v/v) filter-sterilized fetal bovine serum (FBS), 2 mM L-Glutamine, and 10 μg/mL gentamycin. HEp2 cells used in all experiments were used at passage 4-15. Low passage (3–8) human endocervical epithelial E6/E7-immortalized End1s (End1 E6/E7, ATCC CRL-2615) were cultured at 37° C with 5% atmospheric CO_2_ in Keratinocyte Serum-Free Medium (KSFM, Thermo Fisher Scientific) supplemented with human recombinant epidermal growth factor, bovine pituitary extract, 5 mg/mL gentamicin, and 0.4 mM CaCl_2._ McCoy B mouse fibroblasts (originally from Dr. Harlan Caldwell, NIH/NIAID) used for *Chlamydia* propagation were cultured under comparable conditions. All cell lines are tested annually for contamination with the Universal Mycoplasma Detection Kit (ATCC 30-1012K).

### *Chlamydia trachomatis* infections and treatment conditions

*Chlamydia trachomatis* serovar L2 (434/Bu) was propagated in McCoy cells and EBs were purified using a 30% Gastrografin density gradient as described previously (61). For infections, HEp2 cultures at confluence 70-80% were washed with Hanks Buffered Saline Solution (HBSS, Gibco, ThermoFisher Scientific, Cat no.), and *Chlamydia* inoculum prepared in HBSS at the indicated MOI was overlaid onto the cell monolayer. Infection synchronization was done by centrifugation at 500 x *g* for 15 min at 4°C. Plates were then incubated at 37°C for 30 min before replacing the *Chlamydia* inoculum with pre-warmed complete DMEM. Treatment with recombinant 10 U/mL human IFNγ (ThermoFisher Scientific RIFNG100) was done at 24 hours post-infection, unless stated otherwise. To inhibit chlamydial protein synthesis, media was replaced with DMEM supplemented with 50 ng/mL of chloramphenicol at the indicated timepoint and kept until experiment termination.

### Quantitative imaging of STAT1 activation

HEp2 cells were seeded onto 12 mm acid-etched glass coverslips (Chemglass, CLS-1760-012) placed in 24-well plates. Cells were infected with *C. trachomatis* serovar L2 (434/Bu) at an MOI of 2 for 24 hours and left untreated or treated with 10 U/mL of IFNγ (ThermoFisher Scientific RIFNG100) for another 6 hours. Cells were fixed with 4% paraformaldehyde and permeabilized with 1:1 ratio of ice-cold methanol-acetone. Fixed monola were blocked with 4% BSA-PBST and probed with primary antibodies for pSTAT1-Tyr701 (ThermoFisher Scientific MA515071, 1:250 dilution) and *Chlamydia* heat-shock protein 60 (cHsp60) (ThermoFisher Scientific MA3-023, 1:2000 dilution), followed by secondary antibodies AlexaFluor 488 anti-mouse (ThermoFisher Scientific, A28175, 1:400 dilution) and AlexaFluor 594 anti-rabbit (ThermoFisher Scientific A-11012, 1:400). Images were acquired using a Nikon CSU-W1 spinning disk confocal microscope with 0.2 μM z-slices from five different fields per coverslip. Raw images were processed in ImageJ with untreated-uninfected (no IFNγ) group as reference for standardizing image brightness and contrast. For quantifying pSTAT1 fluorescence intensity, ImageJ was used to manually outline cell nuclei in the DAPI (blue) channel of Z-projected images, which were then used to measure nuclear pSTAT1 fluorescence intensity in the red channel.

### RNA isolation and RT-qPCR

HEp2 cells were seeded into 6-well plates and allowed to adhere overnight. Cells were infected with *C. trachomatis* serovar L2 (434/Bu) as stated above at an MOI of 4 to ensure the majority of cells were infected. Cells were terminated at indicated timepoints for time course experiments or treated with 10 U/mL of IFNγ as described above. TRIzol reagent was used to lyse cells and lysates were processed using Ambion PureLink RNA Mini kit (ThermoFisher Scientific 12183018A) and complementary DNA (cDNA) was generated using SuperScript IV VILO Master Mix kit (ThermoFisher Scientific 11756050) according to the manufacturer’s instructions. cDNA was diluted 1:10 and added to PowerUp SYBR Green Master Mix (Thermo Fisher Scientific 4364344) with commercial RT-qPCR primers (KiCqStart SYBR Green Primers, Millipore Sigma KSPQ12012) and was assayed in Applied Biosystems QuantStudio 3 Real Time PCR System with standard cycling conditions. Primers were subjected to dissociation curve analysis to ensure that a single product was generated. Cycle threshold (Ct) values were acquired, and relative transcript expression was quantified using the ΔΔCt method, using HPRT1 for normalization.

### Protein lysate preparation and Western blotting

HEp2 cells were infected and treated as described above. Cells lysates were collected and processed as previously described to ensure inhibition of protease and phosphatase activity (62). Proteins were resolved via SDS-PAGE in 7.5% polyacrylamide gels and transferred to 0.45 μm-pore PVDF membranes (Millipore, IPVH00010) using a semi-dry transfer method. Membranes were blocked with 2% normal goat serum (Sigma-Aldrich G6767), and incubated with primary antibodies overnight at 4 °C: pSTAT1-Tyr701 (ThermoFisher Scientific MA515071, 1:1000 dilution), STAT1 (Santa Cruz Biotechnology SC-417, 1:1000 dilution), pJAK2-Tyr1008 (Cell Signaling Technologies 8082-CST, 1:500 dilution), JAK2 XP (Cell Signaling Technologies 3230-CST, 1:500 dilution), IDO1 (Cell Signaling Technologies 86630, 1:1000 dilution), cHsp60 (ThermoFisher Scientific MA3-023, 1:10,000 dilution), vimentin (Abcam ab8978, 1:1000), and α-tubulin (Abcam ab52866, 1:10,000 dilution). Membranes were washed and labelled with secondary goat anti-rabbit HRP (Dako P0448 1:3000 dilution) or rabbit anti-mouse HRP (Dako P0161 1:3000 dilution). Labelled membranes were developed with Immobilon HRP substrate (Millipore Sigma WBKLS0500) and imaged with an Azure Biosystems c600. Image processing and densitometric analyses were done using ImageJ.

### Graphs and statistical analysis

Bar graphs and violin plots were generated using ggplot2 base package (version 4.2.3) as part of the Tidyverse package (https://cran.r-project.org/web/packages/tidyverse/index.html) in rStudio (version 4.2.2). Welch two sample t-test was used to determine statistical significance with p-values less than 0.05 being considered significant.

---

**Figure S1.**
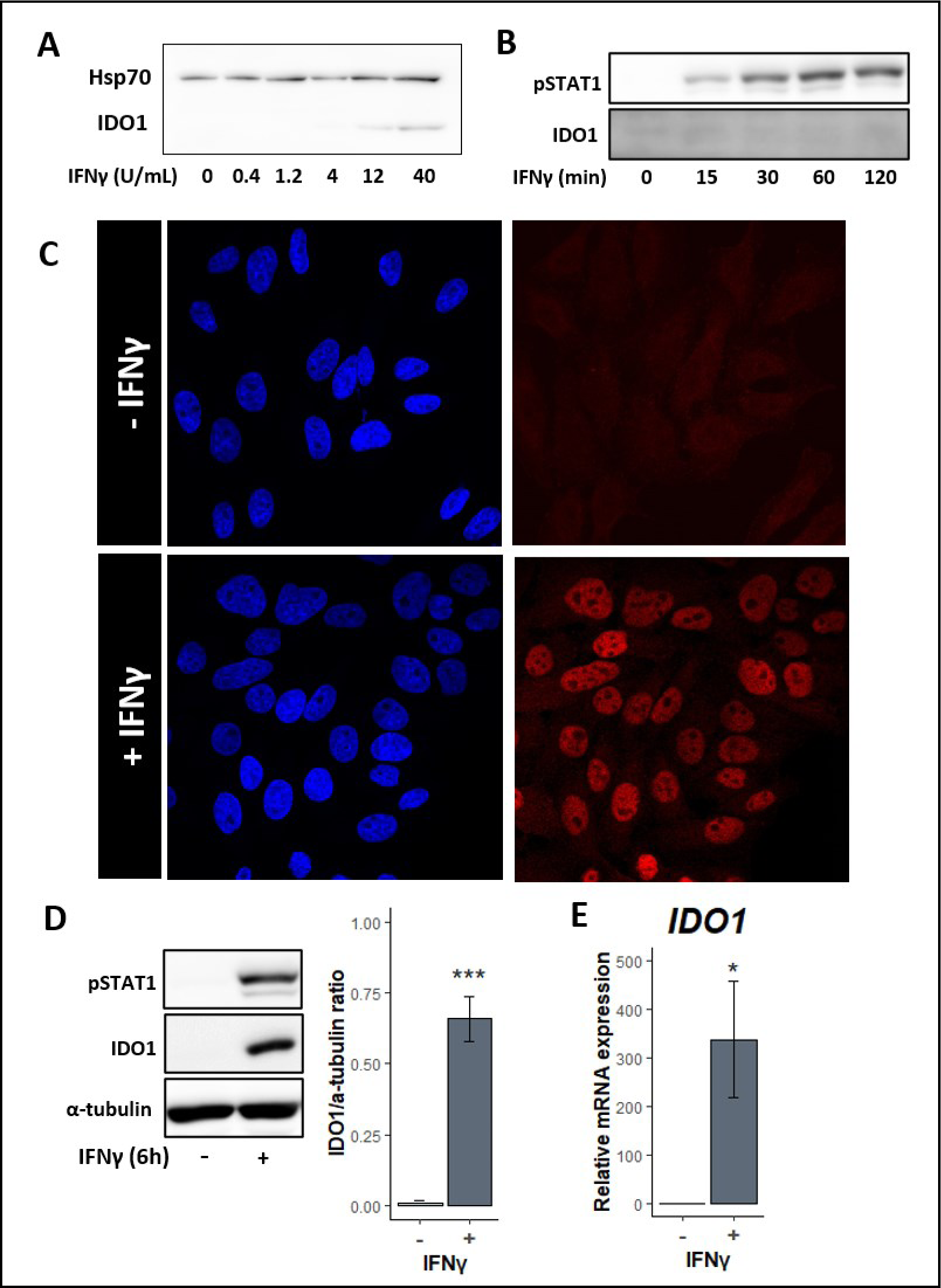
Treatment of Hep2 cells with IFNγ at 10 U/mL is enough to induce activation of STAT1 and expression of the downstream target IDO1. **(A)** Immunoblot analysis of activated STAT1 (pSTAT1) and IDO1 following exposure to 10 U/mL of IFNγ at different exposure lengths. **(B)** Immunofluorescence assay showing nuclear localization of activated STAT1 in the presence of IFNγ **(C)** Immunoblot and densitometric analysis of protein and **(D)** RT-qPCR of transcript expression levels of IDO1 following treatment with IFNγ at 10 U/mL at 24h hpi for 6 hours. All data are representative of 4 independent experiments, and statistical significance was determined using Welch’s t-test. **p < 0.01, *p < 0.05, ns = not significant.

**Figure S2.**
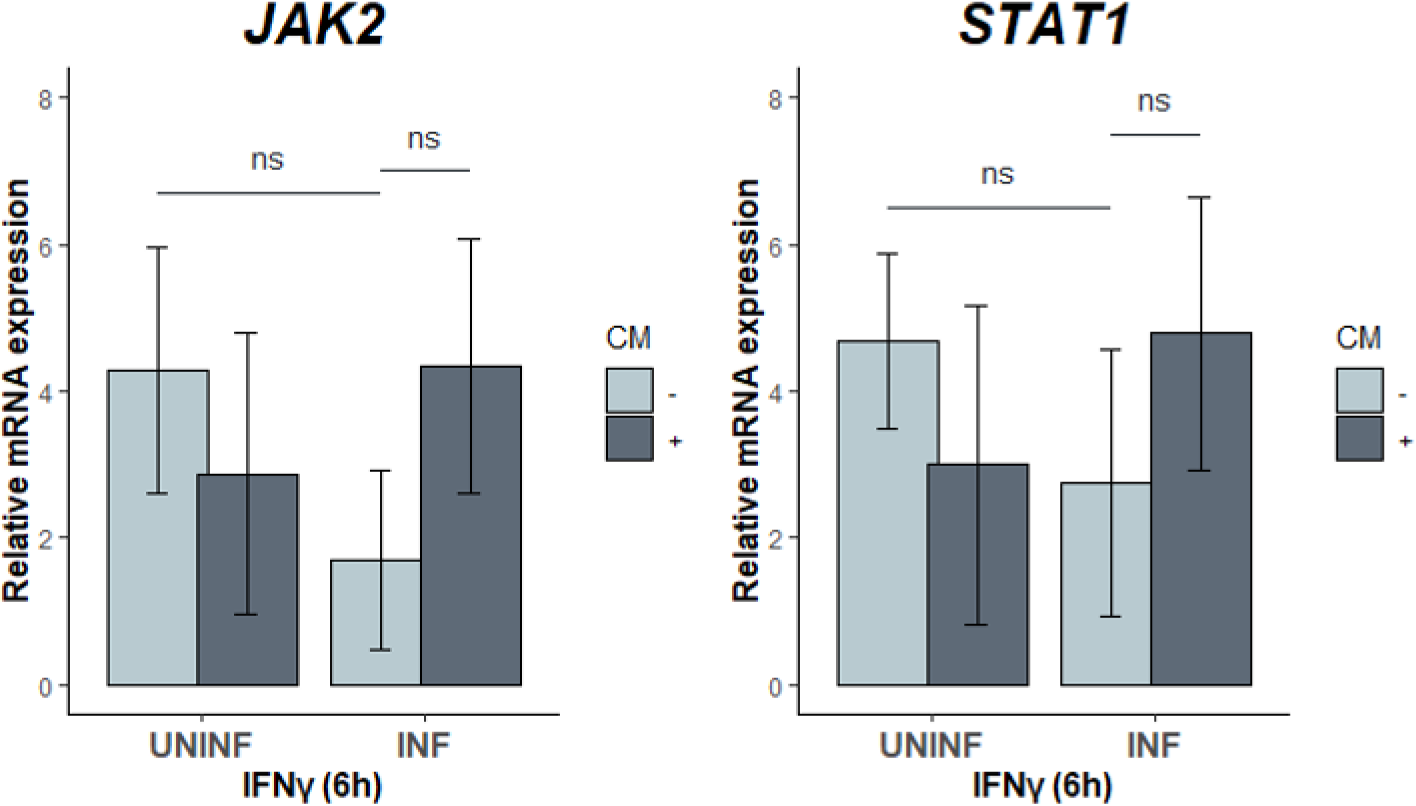
*C. trachomatis* serovar D down-modulates of JAK2 and STAT1 expression in infected HEp2 cells. RT-qPCR analysis of JAK2 and STAT1 transcript expression in HEp2 cells that infected with the genital serovar at MOI=4. Chloramphenicol (CM) was added at 8 hpi, while 20 U/mL IFNγ was added for 6h starting at 24hpi prior to RNA collection. All data are representative of 3 independent experiments, and statistical significance was determined using Welch’s t-test. ns = not significant.

**Figure S3.**
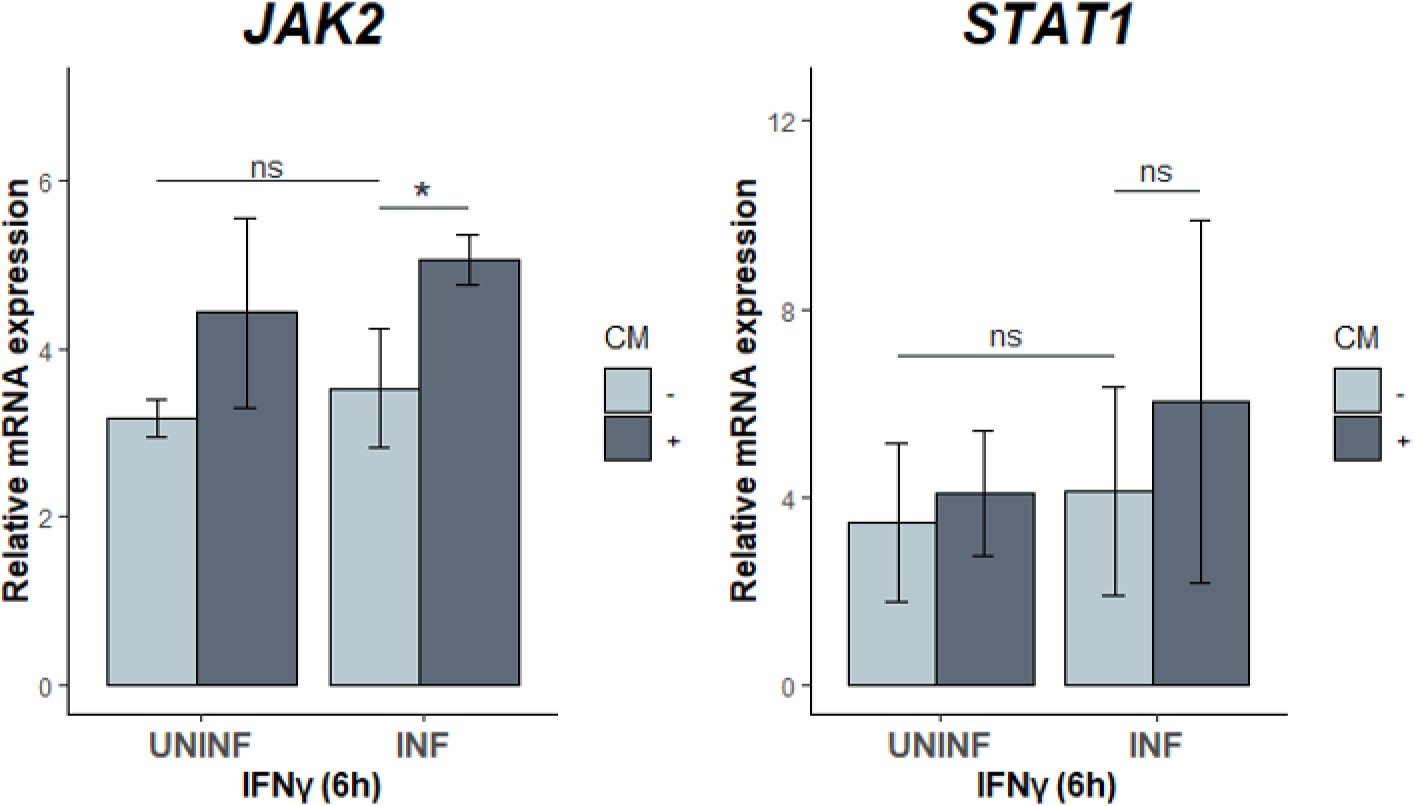
*C. trachomatis* serovar L2 downmodulates expression of JAK2 and STAT1 transcript in infected End1 cells. RT-qPCR analysis of JAK2 and STAT1 transcript expression in endocervical End1 cells that are infected with *C. trachomatis* lymphogranuloma venereum (LGV) serovar L2 at an MOI=4. Chloramphenicol (CM) was added at 8 hpi, while 20 U/mL IFNγ was added for 6h starting at 24hpi prior to RNA collection. All data are representative of 4 independent experiments, and statistical significance was determined using Welch’s t-test. **p < 0.01, *p < 0.05, ns = not significant.

**Figure S4.**
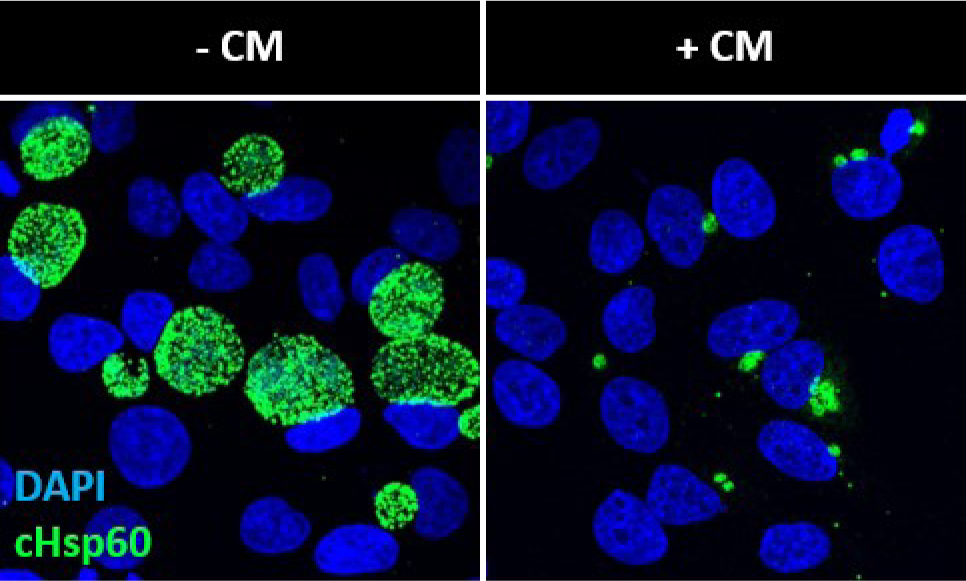
Chloramphenicol treatment limits chlamydial growth. Immunofluorescence visualization of 24hpi *chlamydial* inclusions following chloramphenicol treatment (CM) at 8 hpi observed by staining with chlamydial Hsp60 (cHSP60) antibody in HEp2 cells.

